# Novel co-culture plate enables growth dynamic-based assessment of contact-independent microbial interactions

**DOI:** 10.1101/145615

**Authors:** Thomas J. Moutinho, John C. Panagides, Matthew B. Biggs, Gregory L. Medlock, Glynis L. Kolling, Jason A. Papin

**Affiliations:** Department of Biomedical Engineering, University of Virginia, Charlottesville, VA USA

**Keywords:** phenotypic screening, growth curve, multiwell plate assay, metabolism, laboratory automation, metabolic interactions, dynamic growth measurement, co-culture, contact-independent interaction, contact-dependent interaction, microbial interactions

## Abstract

Interactions between microbes are central to the dynamics of microbial communities. Understanding these interactions is essential for the characterization of communities, yet challenging to accomplish in practice. There are limited available tools for characterizing diffusion-mediated, contact-independent microbial interactions. A practical and widely implemented technique in such characterization involves the simultaneous co-culture of distinct bacterial species and subsequent analysis of relative abundance in the total population. However, distinguishing between species can be logistically challenging. In this paper, we present a low-cost, vertical membrane, co-culture plate to quantify contact-independent interactions between distinct bacterial populations in co-culture via real-time optical density measurements. These measurements can be used to facilitate the analysis of the interaction between microbes that are physically separated by a semipermeable membrane yet able to exchange diffusible molecules. We show that diffusion across the membrane occurs at a sufficient rate to enable effective interaction between physically separate cultures. Two bacterial species commonly found in the cystic fibrotic lung, *Pseudomonas aeruginosa* and *Burkholderia cenocepacia*, were co-cultured to demonstrate how this plate may be implemented to study microbial interactions. We have demonstrated that this novel co-culture device is able to reliably generate real-time measurements of optical density data that can be used to characterize interactions between microbial species.

## Introduction

There exists an extensive amount of interaction among microorganisms in microbial communities [1–3]. An improved understanding of these interactions and their governing mechanisms in a physiologically relevant context will enable more informed treatment of polymicrobial infections and more precise modulation of microbial communities [4–6]. Interactions between microbes are characterized using a variety of methods [7]. Many interactions that take place within microbial communities are due to diffusible molecules such as cross-fed metabolites, quorum sensing molecules, and antimicrobial compounds [8,9]. For example, muricholic acid, a microbe derived secondary bile acid inhibits *Clostridium difficile* taurocholic acid-mediated spore germination [10]. Interactions mediated via diffusible molecules generally do not require the physical interaction of cells and are thus contact-independent [11–14]. These interactions are challenging to characterize with existing approaches [12].

Common co-culture techniques include well mixed co-cultures [15], conditioned media exchange [16], agar plate colony assays [17,18], and membrane divided co-culture such as Corning ® Transwell ® co-culture plates [19]. Each of these methods are limited in their ability to phenotypically characterize the growth dynamics of the microbes in co-culture. In a mixed co-culture it is challenging to measure the individual growth curves of the two species using high-throughput techniques. It is possible to use qPCR techniques to determine the relative abundance of each species; however, this is a technically and logistically challenging experimental technique requiring the development of specific primers for each species [20,21]. Conditioned media exchange experiments are limited to unidirectional interactions which do not capture the dynamic response of cells to changing conditions [16]. The Corning ® Transwell ® culture plates keep cells physically separate while allowing for contact-independent interactions, yet the horizontal membrane does not allow for the collection of optical density based continuous growth curve data for each culture.

Since the advent of semipermeable membrane-divided co-culture tools [22,23], to the best of our knowledge, this concept has never been interfaced with automated plate reader technology for the high-throughput continuous quantification of optical density-based phenotypic assessment of interacting cultures. Optical density of liquid bacterial cultures has been used for a multitude of phenotypic studies that aim to determine the relative changes in cellular growth subject to various environmental conditions [24–29]. We present a novel co-culture plate with a vertically oriented membrane that maintains physical separation of two liquid cultures, yet allows for real-time contact-independent interactions across the membrane. The vertically oriented membrane allows for the co-culture plate to interface with a standard 96-well plate reader that is able to continuously monitor the optical density of both cultures on either side of the membrane. This culture tool is a simple, convenient, and inexpensive method for generating individual growth curves of two batch bacterial cultures as they interact across a membrane.

## Materials and Methods

### Strain Information

We used *Escherichia coli* (K12), *Pseudomonas aeruginosa* (PA14), and *Burkholderia cenocepacia* (K56-2) in this study.

### Media Preparation

Lysogeny broth – Miller (LB) medium: tryptone (10g/L), yeast extract (5g/L), NaCL (10g/L), pH was adjusted to 7.0 with NaOH. In several experiments the LB media was diluted with 1x Dulbecco’s Phosphate Buffered Saline (DPBS) (Gibco by Life Technologies). This dilution is indicated throughout the paper as the percentage of LB that is in the diluted media.

### Sterilization Procedures

Before each experiment all parts of the co-culture plate were steam autoclaved at 121°C, 100kPa, for 60 minutes. The polycarbonate membranes (Isopore^TM^ Membrane Filter, 0.1 µm VCTP; EDM Millipore) were prepared by soaking in 70% ethanol for 10 minutes. For further description and rationale, see S1 Appendix. In a biosafety cabinet, the ethanol-soaked membranes were clamped between the wells and the assembly was left for 10 minutes to allow the ethanol to evaporate. For plate assembly protocol and visual aids, see S1 Appendix.

### Counting of Colony Forming Units (CFUs)

CFUs were counted as previously reported [30]. Briefly, a serial dilution down to 10^−7^ for each of the original cultures was prepared, 10 mL of each dilution was dripped onto LB agar plates and left to dry for roughly 10 minutes. The CFU plates were then incubated for the appropriate amount of time for visible colony growth. Colonies were then manually counted. Reported counts were done in quadruplicate (n=4).

### Growth Curve Collection and Processing

Each well of the co-culture plate was loaded with 2 mL of media. Where appropriate, wells were inoculated at a calculated OD_600_ of 0.0005 with the bacterial strain specified. The co-culture plate was then placed into a Tecan Infinite M200 Pro, incubated at 37°C, shaken linearly at 3mm 450 rpm, and OD measurements were recorded at 600 nm every 5 minutes. All of the experiments were conducted in triplicate with biological replicates. The data from each experiment was exported as an Excel file and processed in MATLAB (R2014b; Mathworks). The growth curve plots consist of the average (bold line) displayed with the maximum and minimum values (as shaded regions around the average line). All growth experiments were conducted in triplicate. The MATLAB scripts used for all data processing are available, refer to the Availability section.

### Scanning Electron Microscopy (SEM)

Following an *E. coli* experiment with the co-culture plate, the polycarbonate membranes were fixed for 30 minutes with glutaraldehyde (2% by vol.). Followed by three 5-minute rinse steps in 1x DPBS. Samples were then dehydrated using increasing concentrations of ethanol, 10 minutes each in 30, 50, 70, 80, 90, 100, 100% (ethanol in water). The membranes were further dehydrated for 10 minutes in HMDS (hexamethyldisilazane; Sigma). Finally, the membranes were stuck to SEM stubs with adhesive carbon strips using the Phenom starter kit (Ted Pella, Redding, CA, USA) and sputter coated with gold using a SCD005 sputter coater (Bal-tec, Los Angeles, CA, USA). The final samples were imaged using a Sigma VP HD Field-emission SEM (Zeiss, Pleasanton, CA, USA) at 10,000x magnification through the University of Virginia Advanced Microscopy Facility.

### Device Design and Machining

All of the parts for the co-culture plate were designed in SolidWorks 2015; all of these files are available, refer to the Availability section. The files were exported as STL files (also available) and G-code was written for CNC machining. The aluminum parts were cut using a waterjet cutter and the holes were tapped by hand. The polypropylene wells were started using a waterjet cutter and finished using a milling machine. The polycarbonate was also cut using the waterjet cutter. All parts are designed to be able to be CNC machined without the use of a waterjet cutter. The silicone gaskets were made using a laser cutter (Universal laser Systems X-660 with a 50 watt CO_2_ laser). For a detailed list of the parts please refer to the S1 Appendix and for part specifications see S1 Technical Drawings.

## Results

### Design and Description

The vertical membrane co-culture plate consists of eight co-culture chambers, each chamber is composed of two wells separated by a membrane that is replaced before each experiment. Each well is designed to hold 2 mL of culture and this 4 mL of total liquid in each chamber. All of the materials used for the body of the plate can be sterilized via autoclave. The outer dimensions and wells on the plate line up with the dimensions and wells of a standard Corning ® 96-well plate, allowing it to interface with any plate reader designed to read 96-well plates. Each of the co-culture wells lines up with two wells on a 96-well plate, allowing for an internal technical replicate to be collected for each well to reduce noise (Fig 1A).

**Fig 1.**
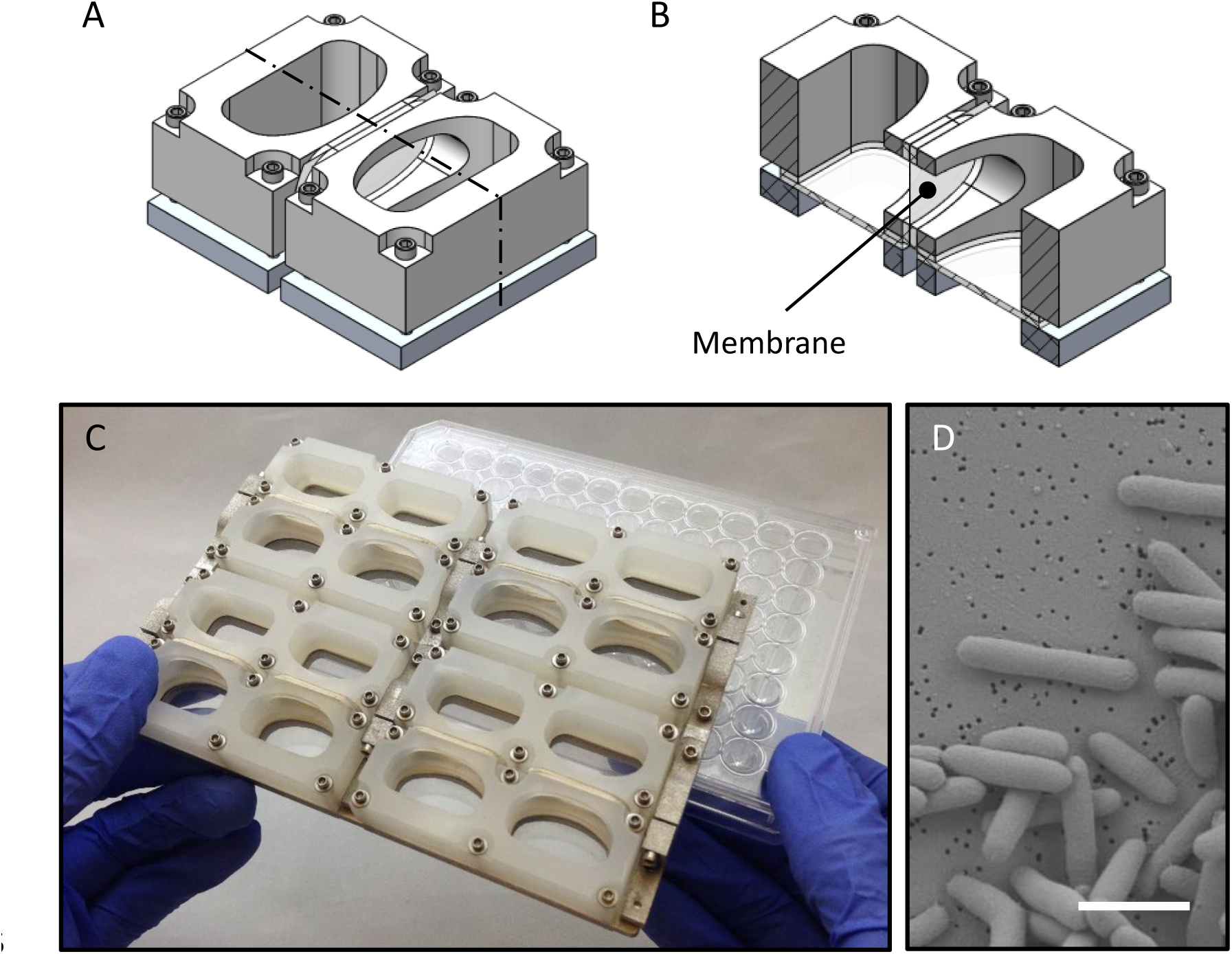
Co-culture plate design. The co-culture plate consists of eight individual co-culture chambers. Each chamber consists of two wells that are able to hold liquid cultures that are physically separated by a semi-permeable membrane that allows for diffusion-mediated interactions. A representative isometric mechanical drawing of a single co-culture chamber is shown in (A), note it is composed of two wells. For a better view of the chamber, a cross-sectional view of it is shown in (B); the semi-permeable membrane is labelled. The co-culture plate composed of eight co-culture chambers has the same profile as a standard 96-well plate. Each well on the co-culture plate aligns with two wells of a 96-well plate and the culture volume is 2 mL per well (4 mL total per chamber) (C). An SEM image captures *E. coli* cells fixed on the surface of a polycarbonate membrane with 0.1 µm pores (D); the scale bar is 2 µm.

The well walls are machined polypropylene, bolted to a machined aluminum base. Clamped between the polypropylene wells and aluminum base are clear polycarbonate pieces acting as the bottom of the wells. A silicone gasket creates a liquid tight seal on the bottom edge of the wells. Additional silicone gaskets are adhered to the side ports in the well walls to create a seal against the membrane which is clamped between the wells. The location of the membrane is indicated in Fig 1B. Any type of membrane can be used in this plate; this point is discussed in the Materials and Methods section.

The base of the plate is composed of three separate parts. Each of the wells is first clamped onto the three base parts and these parts are subsequently clamped together horizontally after the membranes are in place. The dual clamping design allows for adequate force to be applied to create water tight seals both against the bottom of the wells and the sides where the membranes are placed. For further description of the design and machining of the plate, as well as a video of the assembly, refer to the S1 Appendix.

### Validation

The co-culture plate was evaluated for basic functions to guide the interpretation of the data generated using this novel platform. First, we explored whether the rate of metabolite diffusion across the membrane would influence growth dynamics of a culture. Second, we confirmed that the membrane was a sufficient barrier to maintain physical isolation between wells. Finally, we ontrol for later multi-species co-cultures.

We characterized the impact that diffusion of metabolites across the membrane might have on growth characteristics. It was composed of four concentrations of two conditions, ‘pre-mixed’ and ‘gradient’ (Fig 2). The ‘pre-mixed’ condition is inoculated on one side of the membrane and has equal concentrations of LB (diluted with DPBS) on either side. The ‘gradient’ condition is also inoculated on one side of the membrane, but starts with all of the LB on the opposite side. Therefore, for growth to occur on the DPBS-inoculated side, the LB must diffuse across the membrane. The total quantity of LB provided between each condition was held constant. These two conditions were assayed at four different concentrations to demonstrate the observed behavior at various concentration gradients across the membrane, ranges of maximum optical density, and resulting population densities. We observe that there are only slight differences between the paired conditions at all four concentrations. These results indicate that the essential metabolites in LB are able to diffuse across the membrane at a sufficiently rapid rate to allow *E. coli* to grow similarly to the control case.

**Fig 2.**
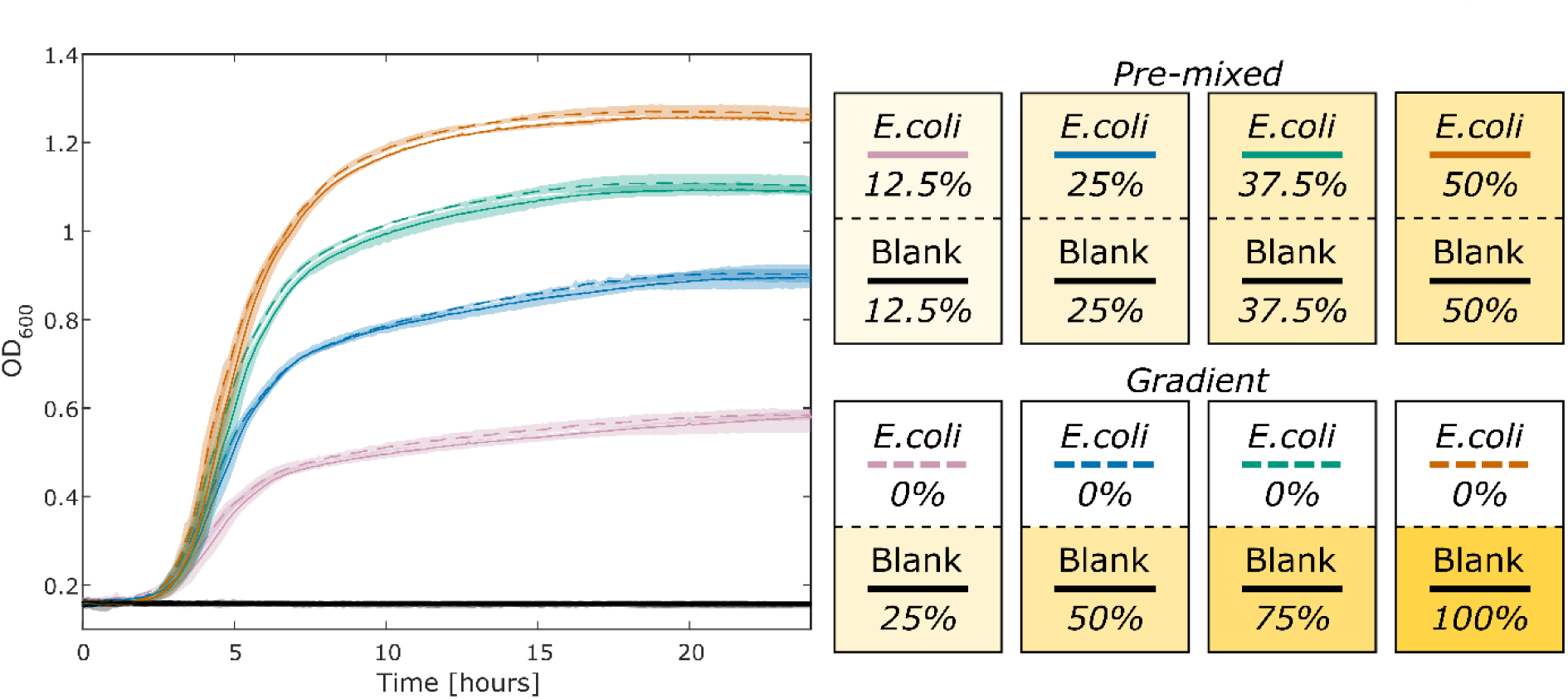
Real-time diffusion of metabolites across a membrane. One side of each co-culture chamber was inoculated with *E. coli*, as seen in the pictorial legend on the right, each box represents a chamber with a black dashed line representing the membrane. The terms ‘pre-mixed’ and ‘gradient’ describe the initial media conditions. The gradient condition was loaded with LB on one side and 1x DPBS on the other. The pre-mixed condition was loaded with LB that was diluted in half with DPBS to simulate complete diffusion of LB across the membrane. These two conditions were tested with four initial concentrations of LB, 25%, 50%, 75%, and 100%, all diluted using 1x DPBS. The final pre-mixed concentration of the medium for each well was 12.5%, 25%, 37.5%, and 50% LB. This experiment was cultured as described in the methods for 24 hours. This experiment was conducted in triplicate (n = 3).

The data for Fig 2 was generated in triplicate such that there were 24 individual co-culture chambers inoculated on one side of the membrane only. Of these 24 individual cases, the optical density of the negative control side was measured to test that the membrane serves as a sufficient barrier to *E. coli* crossing from one well to the other. We never observed *E. coli* contamination from one well to another, therefore the design of the plate and size of the pores in the membrane (Fig 1D) are sufficient to maintain complete physical separation between the two sides of each chamber and yet allow for the exchange of nutrients and small molecules to support growth without a notable defect in the associated growth dynamics for the conditions we tested. Pictures of the plate can be seen with co-cultures at the end point in the Supporting information (Fig S1).

One potential application for this co-culture plate is the characterization of growth dynamics for two different species on either side of the membrane. To determine the basic characteristics of co-culture between competing cultures, we cultured *E. coli* in isolation on one side of the chamber for one condition and two *E. coli* populations were cultured in adjacent wells separated by the membrane and thus competing for nutrients (Fig 3). Both of these conditions were assessed at 50% LB (diluted with 1x DPBS) and 100% LB. The condition in which *E. coli* is isolated on just one side of the co-culture chamber acts as a reference point compared to the case in which two *E. coli* populations are competing. For the condition in which *E. coli* is competing and cultured with 100% LB, the growth characteristics are similar to those observed when *E. coli* is isolated and cultured in 50% LB. These data indicate that the isolated and competing conditions with a certain microbe and media condition can act as an approximation for the hypothetical case in which two different species in co-culture on either side are in complete metabolic competition with each other. In this context, complete metabolic competition means that the cultures on either side have the same metabolic requirements, this is only the case when the same species are on both sides of the membrane. The growth curve representing complete metabolic competition can be used in tandem with the isolated condition in which there is no metabolic competition for a phenotypic assessment of interactions between two different species in co-culture across the membrane.

**Fig 3.**
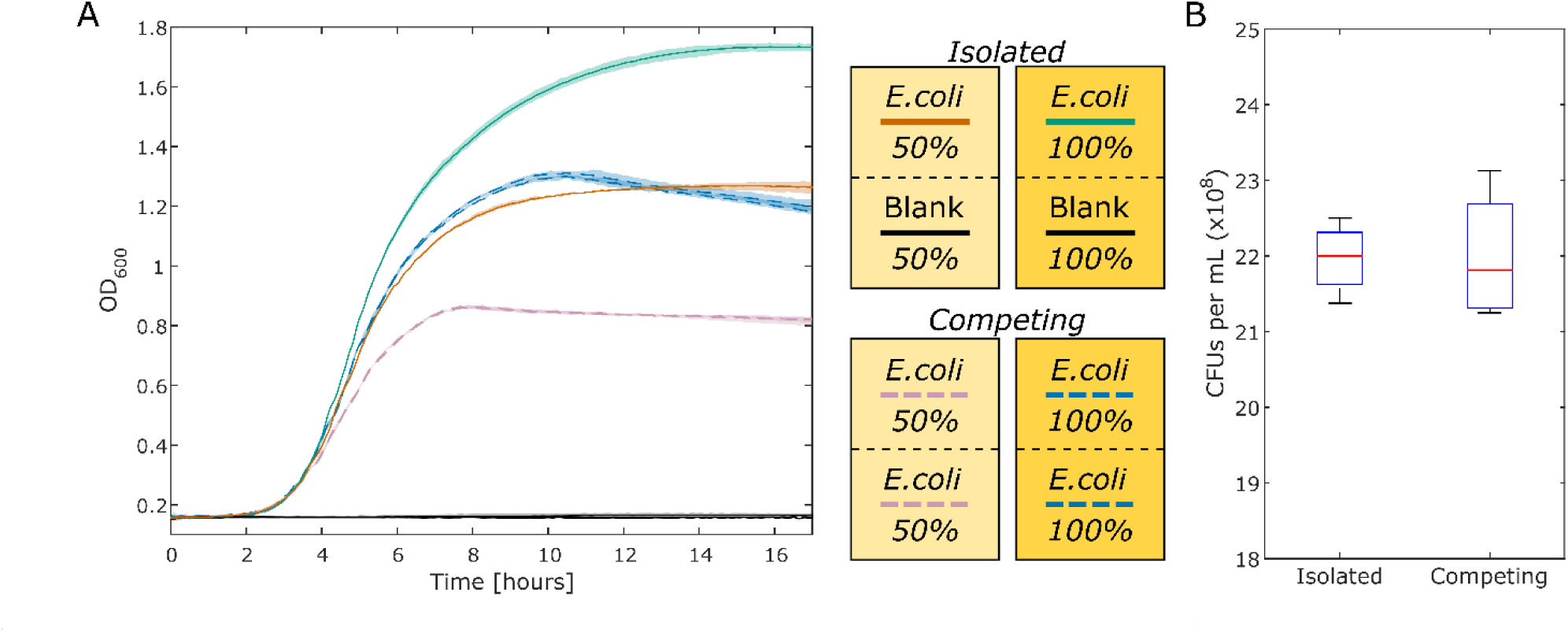
Comparison of isolated versus competing cultures. (A) The green (100% LB) and red (50% LB) lines are the isolated culture condition that have *E. coli* cultured on only one side of the membrane with blank media on the other. The OD for the side of the well that is not inoculated is plotted in black (it maintains the original OD; there is no growth, as expected). In this condition, *E. coli* has access to all of the nutrients on both sides of the membrane, but cell growth is physically constrained to one side. The blue (100% LB) and purple (50% LB) dashed lines are the competing culture condition that have *E. coli* cultured on both sides of the membrane. For the competing cultures, the growth curves from both sides are plotted individually. In this condition, each *E. coli* population must compete for the available nutrients. The maximum and minimum values of the generated growth curves, conducted in triplicate, are displayed as shaded regions around the plotted averages. (B) The biomass produced is approximated by the CFU count of each culture. The CFU counts for the isolated condition as displayed are divided in half to compare to the competing condition, discussed further in the text. These data are the result of four experiments. The boxplot whiskers represent +/- 2.7σ from the mean.

A follow up experiment was conducted to determine if the same number of CFUs, from both sides of the membrane, are produced in the competing versus the isolated conditions. Samples were taken from the 50% LB isolated and competing conditions at 10 hours into incubation. The 50% LB condition was chosen to limit the impact of OD non-linearity and inhibition of growth due to spatial restrictions and prioritize nutrient depletion as the major limiting factor on biomass production. The bacteria were diluted, plated, and CFUs were counted (Fig 3B). The CFU counts for the isolated condition were divided in half to adequately compare to one side of the competing condition. This was done because all of the biomass in the insolated case is located on one side whereas the biomass is split evenly on either side in the competing condition. It can be seen that the same total number of CFUs are present in both the competing and the isolated conditions from Fig 3A. The equivalence between the two conditions in this boxplot indicates that the same number of viable cells are produced in the two different conditions.

### Co-culture of Multiple Species

Infection with *P. aeruginosa* (PA) is pervasive in cystic fibrosis patients [31]. Co-infection with *B. cenocepacia* (BC) can lead to increased mortality rates [32]. These pathogens have been shown to interact in cystic fibrosis infections [33]. We used the co-culture plate to determine the growth characteristics when PA and BC in which media, nutrients, and small molecules are shared. The condition in which a microbe is competing with itself across the membrane, is an approximation of complete metabolic competition. The competing and isolated conditions, as defined in Fig 3, can be used as points of reference when assessing the impact another species has on a culture. In this case, we can see that when PA and BC are co-cultured, BC growth is negatively impacted by the presence of PA (dashed purple), more so than when it is competing with itself (dashed red). However, it appears that PA is unaffected by the present of BC (solid purple vs. solid blue).

## Discussion

In this study, we present a novel tool to enable dynamic growth measurements of individual species interacting in co-culture. Mixed co-culture studies rely on a number of methods for differentiating between specific species when a semi-permeable barrier is not utilized. When applied to mixed co-culture experiments, CFU assays require that populations can be discriminated based on colony morphology [34]. Similarly, flow cytometry based counting assays require discrimination by cellular morphology [35]. Neither of these assays can be used to study co-cultures of morphologically similar populations. Species-specific qPCR assays can be used when genomic sequences are available [20,36,37]. However, manual sampling requires sufficient volume for DNA extraction and therefore greatly constrains possible experimental designs. This requirement of large culture volumes is a limitation shared by all methods that require periodic manual or automated sampling of the culture. Additionally, RT-qPCR assays must be developed for each species in a co-culture study while limiting nonspecific amplification. Species-specific delivery or expression of fluorescent markers are used to discriminate between microbes [34], but several experiments are required during the design of each marker to ensure specificity and stability of the marker. Additionally, genetic alteration of the microbes of interest may be undesirable. While most of these methods can be used in a broader context than batch co-culture in a liquid medium, the experimental design and optimization required for them limits throughput relative to the co-culture plate presented here.

As presented, this novel co-culture plate is able to maintain physical separation of two interacting cultures, while allowing for diffusion mediated interactions. Metabolites across the membrane appear to diffuse across at a sufficiently high rate to not be a limiting factor for growth dynamics. We are able to use the plate to investigate the co-culture of two different species with the use of self-competing controls and isolated culture controls. These controls can be used as a reference for the experimental condition of two species interacting across the membrane.

A particular strength of this co-culture plate is the ability to measure optical density data in real-time. This high temporal resolution captures complex growth dynamics that might not be observed with methods that require manual sampling of the culture. Separating the microbial cultures with a membrane eliminates the need to differentiate the individual species in co-culture. Furthermore, no genetic tools are required in order to screen microbes in this co-culture plate. One possible use of this device could be to co-culture a single species in one well with a complex community in the other well. Although it is not developed here, the wells on the co-culture plate have an adequate volume of media to allow for additional multi-omics analyses to be conducted on the cultures at the end point of the experiment. Such analyses might involve the evaluation of concentration gradients of metabolites across the membrane, or to conduct transcriptomics of cells that are interacting with each other. Furthermore, this novel tool makes it exceptionally simple to generate phenotypic data on the dynamic interactions between two microbial species. The setup for such an experiment (e.g. Fig 4) requires less than two hours.

**Fig 4.**
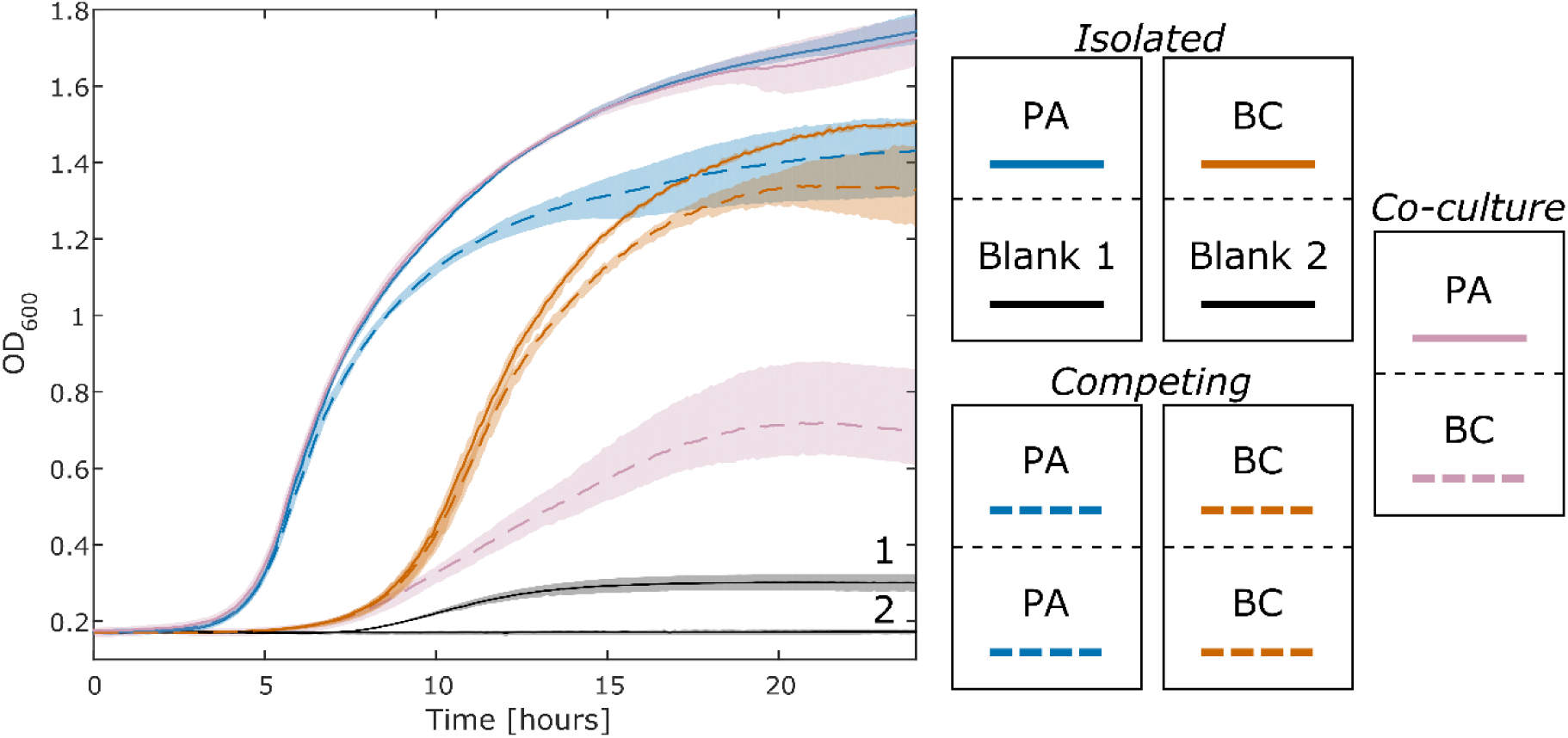
Growth curves of *P. aeruginosa* (PA) and *B. cenocepacia* (BC) in co-culture. The culture conditions can be seen in the legend on the right side of this figure. All wells are started with 100% LB. The purple lines are gathered from the co-culture of PA and BC. The isolated PA and BC cultures are the solid blue and red lines respectively. The black lines are controls from the side of the wells that were not inoculated for the isolated PA and BC cultures. The black line slightly increases (Blank 1) as a result of pyoverdine (produced by PA) that partially absorbs at 600 nm. This result is discussed further in the Supporting information (Fig S2). The growth curves from each of the two competing PA and BC cultures (dashed blue and red lines respectively) are nearly identical (similar to blue and purple in Fig 3) and thus are averaged to simplify the plot. The growth of BC (dashed purple) is negatively impacted when in co-culture with PA (solid purple).

Although the proposed co-culture plate, in its current form, accommodates only one complete two species interaction experiment, throughput can be improved in two ways. Parallelized experiments using additional co-culture plates in conjunction with miniaturized plate readers [38] allows for the collection of endpoint metabolomics samples. As for experiments that do not require such culture volumes, the current co-culture plate design could be scaled down to a format with a greater number of smaller wells. This redesign would be optimized for rapid assays to identify biologically interesting pairs. Additional limitations of the proposed co-culture plate include the restriction to batch culture experiments, and the lack of being able to assess contact-dependent interactions due to physical separation with the membrane.

We have presented a novel co-culture plate that utilizes a vertical membrane to maintain physical separation between two cultures, yet allows for contact-independent interactions. This culture plate allows for high-throughput and high-resolution phenotypic assessment of microbial interactions. As well as interfacing with currently available plate readers, thus allowing for the rapid generation of optical density growth curves.

### Availability

The SolidWorks ® files, STL files, code, and raw data are available at: https://github.com/csbl/CoculturePlate

## Acknowledgements

Nick Anselmo for generating the CNC Machine G-code and machining the parts for the co-culture plate.

## Supporting information

**S1 Appendix. Additional information about the manufacture and use of the co-culture plate.** This document contains a parts list for the plate, a detailed explanation of the design of the plate, and a protocol for using the plate.

**S1 Fig. Endpoint image of co-culture plate after representative experiment from Fig 2.** Note that the lower wells are all void of bacterial growth, while the well on the other side of the membrane is inoculated with an active culture of *E. coli*. Sterility of the wells is maintained by the membranes.

**S2 Fig. Co-culture of *P. aeruginosa* and B. Cenocepacia.** Well 1a is PA, 1b is BC, 5a and 5b are a technical replicate of that, these are the experimental co-cultures. Wells 3a and 3b are the isolated condition of PA, 7a and 7b are the competing condition. Wells 4a and 4b are the isolated condition of BC, 8a and 8b are the competing condition. Wells 2a, 2b, 6a, and 6b are the isolated and competing conditions for PA and BC mixed, these data are not presented in the manuscript. *P. aeruginosa* shows clear production of pyoverdine (green pigment) in the chambers it’s cultured in. The production of pyoverdine has been previously reported (39).

**S1 Technical Drawings. A document containing the technical drawings for each part.** Although these drawings have the specifications for machining the parts, it is recommended that each part is machining based on the CAD files using a CNC machine.

